# A patient-derived *LMX1B* variant causes tissue-specific manifestations of nail-patella syndrome in mice

**DOI:** 10.64898/2026.08.28.747744

**Authors:** Takanori Amano, Keisuke Yoshida, Farzana Sultana, Chigusa Imura, Mayu Shiokawa, Hirotoshi Shibuya, Keishin Takemura, Kyoko Ikeda, Chieko Otsuka, Kazuya Shinbo, Masaru Tamura, Seiya Mizuno

**Affiliations:** Next Generation Human Disease Model Research Team, RIKEN BioResource Research Center, Tsukuba, Ibaraki 305-0074, Japan; Institute for Advanced Medical Sciences, Nippon Medical School, 1-25-16, Nezu, Bunkyo-ku, Tokyo, 113-0031, Japan; Drug Discovery Genetically Modified Animal Platform Unit, RIKEN BioResource Research Center, Tsukuba, Ibaraki 305-0074, Japan; Mouse Phenomics Division, RIKEN BioResource Research Center, Tsukuba, Ibaraki 305-0074, Japan; Laboratory Animal Resource Center in Transborder Medical Research Center, Institute of Medicine, University of Tsukuba, 1-1-1 Tennodai, Tsukuba, Ibaraki, 305-8575, Japan

## Abstract

Nail-patella syndrome (NPS) is a multisystem disorder caused by pathogenic variants in *LMX1B* and is characterized by dysplasia of the nails and patellae as well as extraskeletal complications such as progressive nephropathy and glaucoma. We generated a CRISPR/Cas9 knock-in mouse carrying the R252Q substitution, corresponding to a human *LMX1B* variant associated with renal-predominant disease. Phenotypic analysis revealed that homozygous mice were viable, but they displayed marked growth retardation and severe bilateral ocular opacity. Interestingly, while this model exhibited clear skeletal and ocular defects, the renal phenotype was relatively mild, although increased urinary albumin excretion, focal glomerular basement membrane abnormalities, and subtle changes in renal gene expression were detected. Beyond the classical NPS hallmarks, mutant mice also displayed midbrain morphological abnormalities, suggesting broader developmental consequences of this *LMX1B* variant. This patient-derived variant model not only recapitulates the pleiotropic features of NPS but also demonstrates organ-specific susceptibility to the R252Q substitution, providing a foundation for elucidating the complex molecular mechanisms underlying multisystem disease.

**Author Summary:** Predicting how a disease-associated variant will affect different organs remains a major challenge. Variants in the same gene can produce different combinations and severities of symptoms. We studied a particular variant of *LMX1B* that is predominantly associated with kidney disease in patients by generating mice with the corresponding variant in their *Lmx1b* gene. Homozygous mice developed prominent abnormalities of the eyes, kneecap, and midbrain, whereas kidney involvement was comparatively mild. Thus, the mice displayed several features of nail-patella syndrome, although the relative involvement of different organs differed from the kidney-predominant presentation reported in patients carrying this variant. We also found that the altered LMX1B protein retained partial gene-regulatory activity rather than being completely inactive. Together, our findings show that the effects of an LMX1B variant cannot be predicted from its residual activity alone. Instead, the susceptibility of each organ may depend on the cellular and genetic context in which the variant acts.

## Introduction

Nail-patella syndrome (NPS; MIM #161200) is an autosomal dominant disorder classically defined by a tetrad of nail dysplasia, hypoplastic or absent patellae, elbow dysplasia, and iliac horn formation [1]. Approximately 30-50% of affected individuals develop nephropathy, which may progress to end-stage kidney disease, and some also develop glaucoma [1–3].

NPS is primarily caused by mutations in *LMX1B*, which encodes a LIM-homeodomain transcription factor with pleiotropic roles in embryonic development and adult organ homeostasis [4, 5]. To date, a large number of disease-associated *LMX1B* variants have been catalogued across clinical databases [6, 7]. These pathogenic variants are frequently found in the N-terminal LIM domains, which mediate protein-protein interactions, and in the homeodomain, which confers sequence-specific DNA binding [2]. Pathogenic variants in the homeodomain, especially missense mutations, are frequently associated with renal involvement, presumably by impairing the transcriptional regulation of downstream targets essential for podocyte integrity and glomerular basement membrane homeostasis, such as *NPHS2, COL4A3*, and *COL4A4* [8, 9].

Despite the shared genetic etiology, clinical manifestations show remarkable variability. For instance, a recent report of a family carrying the *LMX1B^S242del^*variant described limb manifestations in monozygotic twins and proteinuria in their affected mother, illustrating that the organ manifestations associated with a given *LMX1B* variant may differ even within the same family [10]. Studies using *Lmx1b*-deficient mice have established its essential roles in specifying dorsal cell fate in the limbs [5, 11], maintaining glomerular podocyte differentiation [8], ocular anterior segment formation [12], and midbrain-hindbrain patterning via the regulation of *Fgf8* and *Wnt1* [11].

While these *Lmx1b* knockout models have provided foundational insights, they face a fundamental translational limitation: homozygous null mice exhibit perinatal lethality, whereas heterozygous null mice often fail to recapitulate the progressive renal and ocular decline observed in human patients [13]. The *Lmx1b^Icst^* mouse model, which harbors a V271D substitution arising from ENU-induced mutagenesis, represented a significant advance by demonstrating a dominant phenotype including glaucoma and semi-lethality driven by interference with LDB1-mediated dimerization [13]. However, because the V271D variant arose from chemical mutagenesis in mice rather than identified in human patients, there remains a need for physiologically relevant animal models carrying patient-derived variants at the corresponding residues.

Among the pathogenic *LMX1B* variants reported to date, the R246Q substitution is clinically notable as a recurrent renal-predominant variant rather than as a variant associated with the classic NPS phenotype. R246Q has been reported in patients with nail-patella-like renal disease (NPLRD) and familial focal segmental glomerulosclerosis (FSGS), in some cases without overt extrarenal manifestations [14–17]. The affected residue is highly conserved across vertebrate species and located within helix II of the LMX1B homeodomain. Consistent with this structural context, functional analysis demonstrated that R246Q partially impairs LMX1B-dependent transcriptional activity [16]. However, it remains unclear whether the renal-predominant phenotype associated with this variant reflects an intrinsic, species-conserved effect of altered LMX1B function or species-dependent differences in tissue susceptibility.

In this study, we generated and characterized a knock-in mouse model harboring the *Lmx1b^R252Q^* substitution, which corresponds to the human *LMX1B^R246Q^* variant, to determine whether its renal-predominant phenotype is recapitulated *in vivo*. Although the mutant mice exhibited prominent skeletal, ocular and brain abnormalities alongside a comparatively mild renal phenotype, this phenotypic profile provides a valuable *in vivo* platform for investigating tissue-dependent susceptibility and the complex molecular mechanisms underlying *LMX1B* pleiotropy.

## Results

### Generation of *Lmx1b^R252Q^* knock-in mice

To investigate the *in vivo* pathogenesis of the NPLRD-associated *LMX1B^R246Q^* variant, we generated a genome-edited mouse model carrying the equivalent substitution at the mouse *Lmx1b* locus (Fig 1A). Sequence analysis confirmed precise introduction of the intended nucleotide substitution without any unexpected insertions or deletions (Fig 1B). Intercrossing of heterozygous mice successfully yielded wild-type, heterozygous, and homozygous offspring. Although *Lmx1b*^R252Q/R252Q^ mice were viable, they displayed growth retardation and small bodies (Fig 1C). Both male and female homozygous mutants exhibited substantially lower body weights compared with wild-type littermates (Fig 1D, E). In addition to the small body size, homozygous mice consistently presented with severe bilateral ocular opacity (Fig 2A, B).

**Fig 1.**
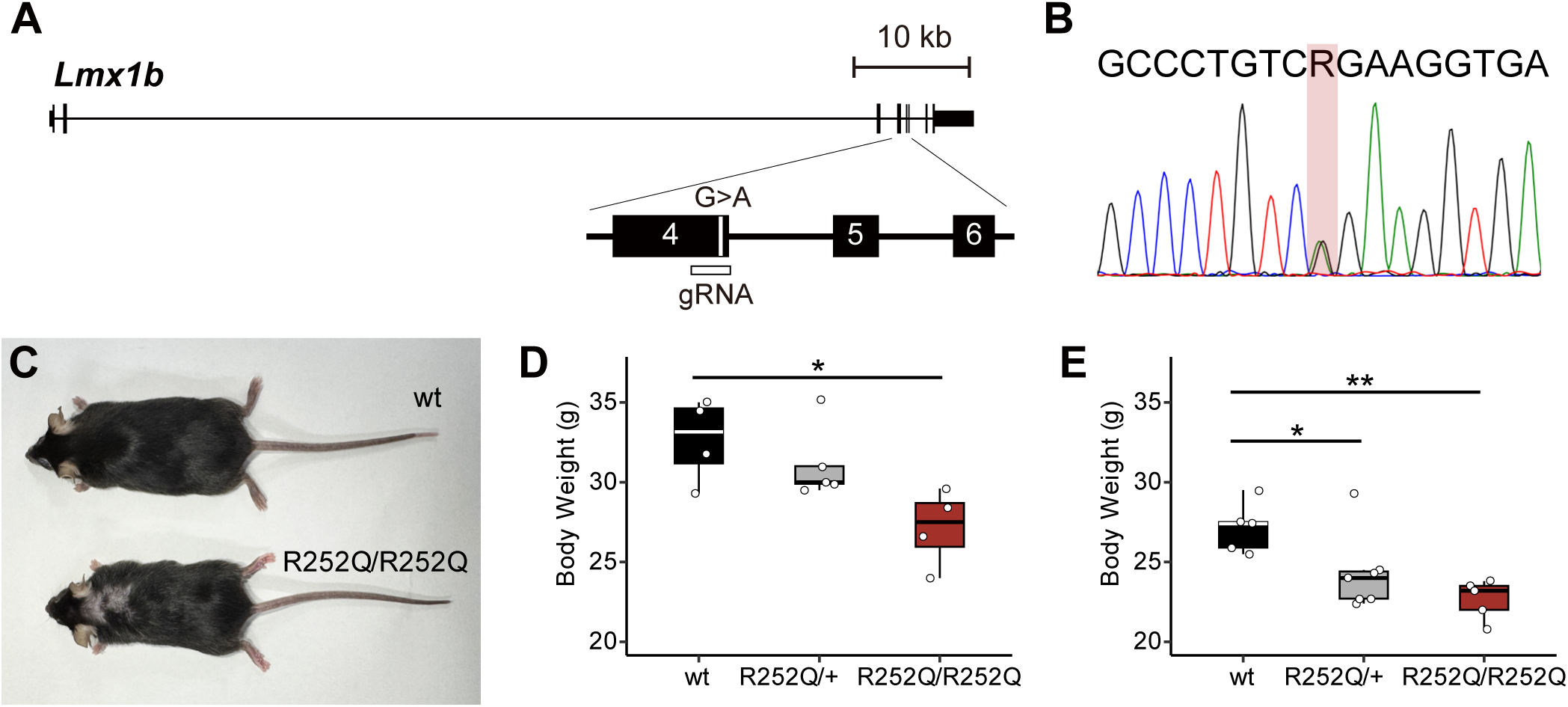
Generation of *Lmx1b^R252Q^*knock-in mice. (A) Schematic representation of the mouse *Lmx1b* locus showing the CRISPR/Cas9 target site and the single-nucleotide substitution introduced to generate the R252Q allele. (B) A representative Sanger sequencing chromatogram of an *Lmx1b^R252Q/+^* heterozygous knock-in mouse confirming the intended nucleotide substitution at the *Lmx1b* locus. (C) Dorsal view of wild-type and *Lmx1b^R252Q/R252Q^* mice. (D, E) Box plots showing body weights of male (D) and female (E) wild-type, *Lmx1b^R252Q/+^*, and *Lmx1b^R252Q/R252Q^* mice at 21 weeks of age. Homozygous mutant mice showed reduced body weight in both sexes. Each dot represents an individual mouse. Statistical comparisons were performed using one-way ANOVA followed by Dunnett’s multiple-comparisons test, with wild-type mice as the control group. \**P* < 0.05; \*\**P* < 0.01.

**Fig 2.**
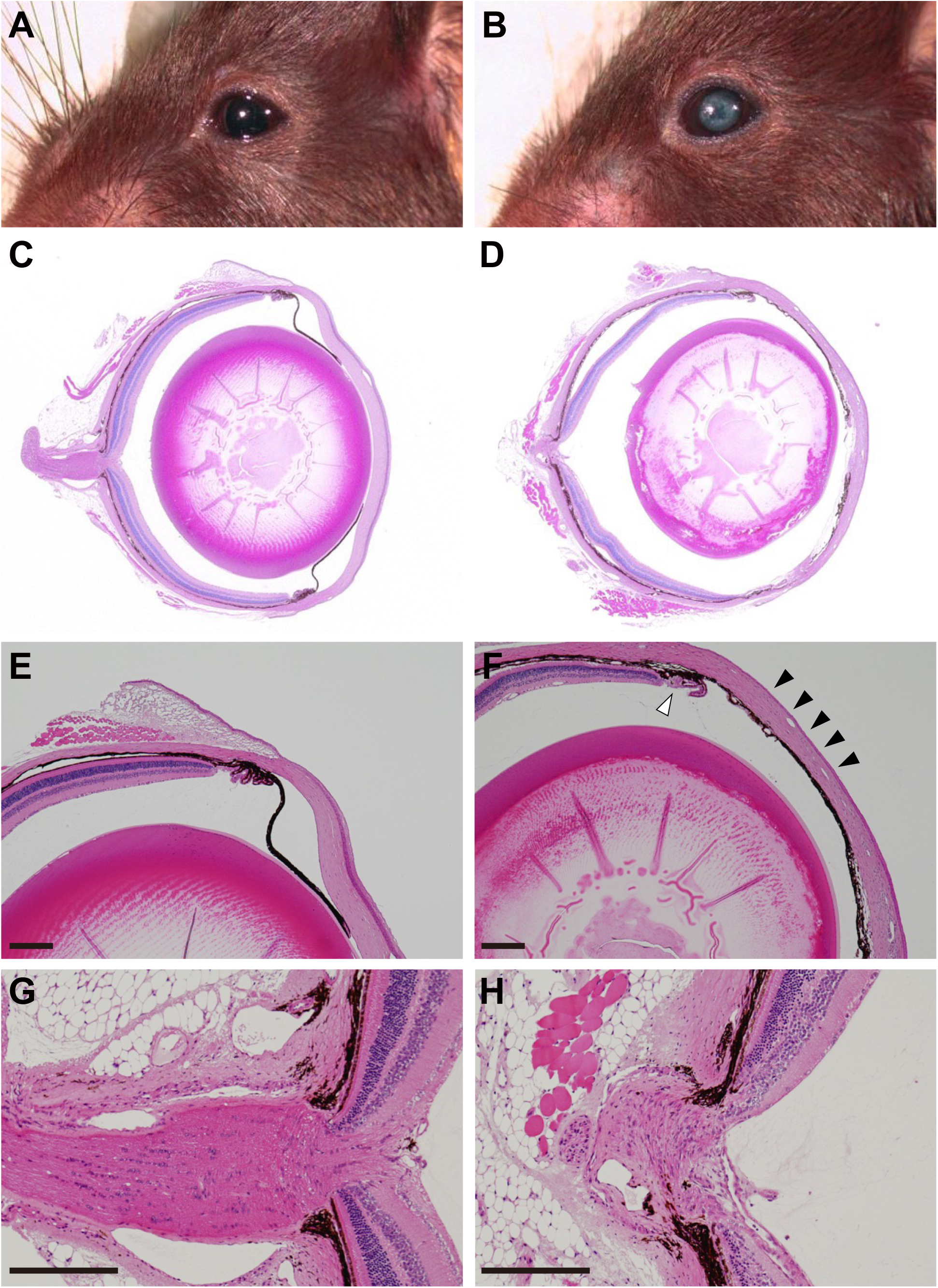
Glaucoma-like ocular pathology in *Lmx1b^R252Q/R252Q^* mice. (A, B) Gross appearance of eyes from wild-type (A) and *Lmx1b^R252Q/R252Q^* mice (B). Homozygous mutant mice showed severe ocular opacity. (C, D) Low-magnification images of HE-stained whole eye sections from wild-type (C) and *Lmx1b^R252Q/R252Q^* mice (D). Homozygous mutant eyes showed abnormal ocular architecture, including thinning of the cornea and retina. (E, F) Higher magnification images of the anterior segment. The mutant eye showed ciliary body hypoplasia (open arrowhead) and adhesion of the iris to the peripheral cornea (arrowheads), with focal closure of the iridocorneal angle. (G, H) Higher magnification images of the optic nerve head from wild-type (G) and homozygous mutant mice (H), showing marked deformation of the optic nerve head in the mutant. Scale bars, 200 µm (E-H).

### Glaucoma-like ocular pathology in *Lmx1b^R252Q/R252Q^* mice

Ocular abnormalities are part of the broader NPS phenotypic spectrum. We, therefore, examined the ocular phenotype of *Lmx1b^R252Q/R252Q^*mice in more detail. Histological analysis revealed marked structural abnormalities in mutant eyes compared with wild-type controls. Low-magnification sections showed thinning of the cornea and retina, and an altered configuration of the anterior chamber in homozygous mutant eyes (Fig 2C, D). In the iridocorneal angle and ciliary body region, mutant eyes exhibited ciliary body hypoplasia and adhesion of the iris to the peripheral cornea, consistent with peripheral anterior synechia. This adhesion was accompanied by distortion and focal closure of the iridocorneal angle (Fig 2E, F). In addition, mutant eyes exhibited marked deformation of the optic nerve head, whereas wild-type eyes showed a well-organized optic nerve head structure (Fig 2G, H). Together with gross ocular opacity, these histological abnormalities are consistent with glaucoma-like ocular pathology in *Lmx1b^R252Q/R252Q^* mice.

### *Lmx1b^R252Q/R252Q^* mice exhibit NPS-associated patellar hypoplasia

Although skeletal abnormalities have not been reported in patients carrying the *LMX1B^R246Q^* variant, we also assessed the limb skeleton in this model, given the limb defects characteristic of NPS and the established role of *Lmx1b* in dorsal limb development. Micro-computed tomography (µCT) of the hindlimb skeleton was performed followed by three-dimensional reconstruction (Fig 3). Whole hind limb reconstructions showed a well-formed ovoid patella at the knee joint, whereas the patella was markedly smaller and altered in shape in *Lmx1b^R252Q/R252Q^* (Fig 3A-D). In some homozygous mice, the hypoplastic patella also appeared displaced relative to the femoral trochlea. Quantitative analysis confirmed significant reductions in both patellar length and volume in homozygous mice (Fig 3E, F). These findings indicate that the R252Q substitution impairs normal patellar development *in vivo*. Despite the clear patellar phenotype, no obvious nail abnormalities were observed on gross examination.

**Fig 3.**
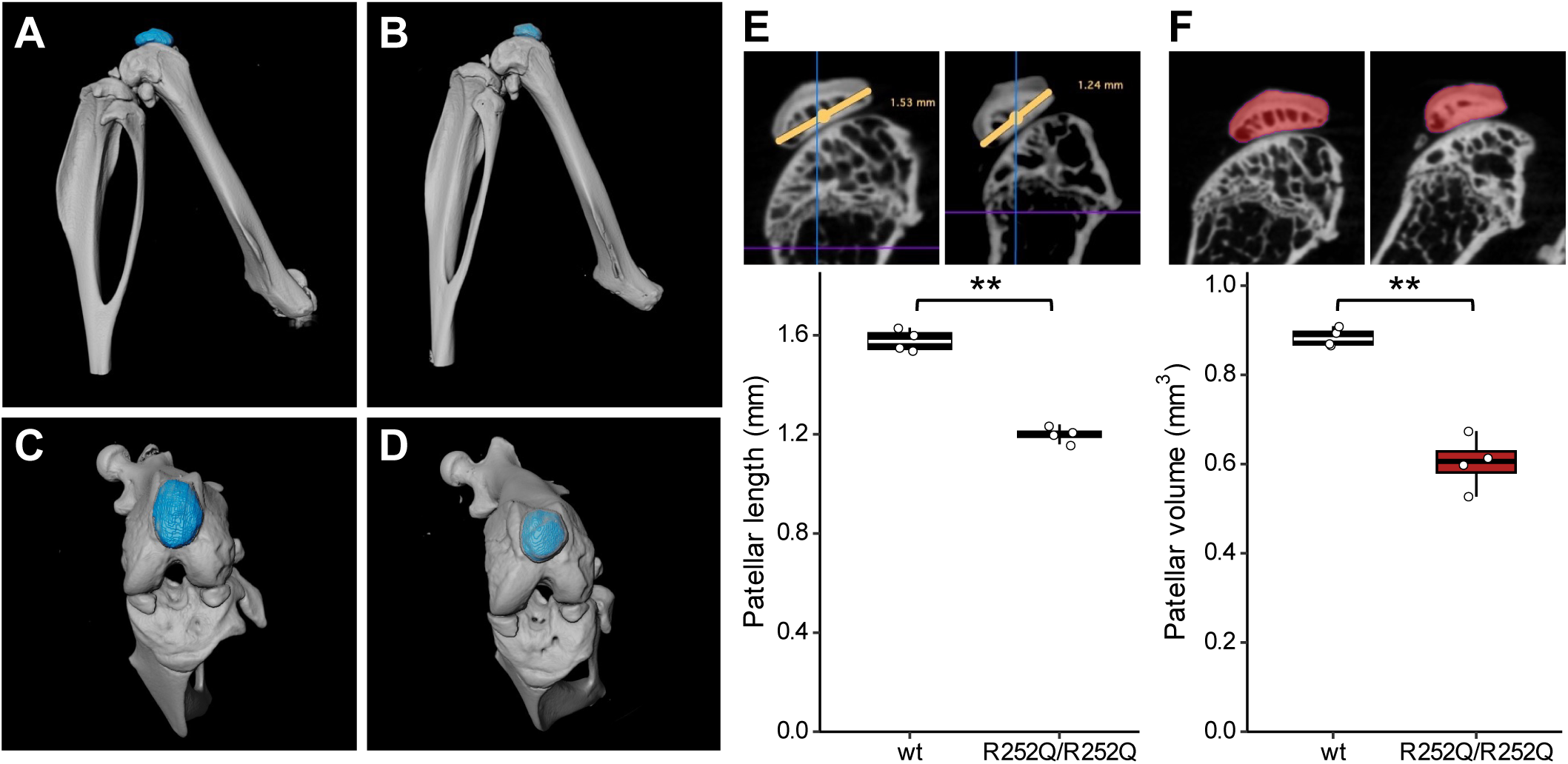
Patellar hypoplasia in *Lmx1b^R252Q/R252Q^* mice. (A-D) Lateral and dorsal views of representative three-dimensional µCT reconstructions of the patellar region from wild-type (A, C) and *Lmx1b^R252Q/R252Q^* mice (B, D). The patellae are highlighted in blue. Homozygous mutant mice showed a reduced and dysmorphic patella compared with wild-type controls. (E) Representative virtual sections showing measurement of the patellar long-axis length (yellow lines), with quantification shown below. (F) Representative virtual sections showing segmentation of the patellar bone (red), with quantification shown below. Patellar bone volume was calculated by integrating the segmented bone regions across serial virtual sections. Both patellar length and volume were significantly decreased in *Lmx1b^R252Q/R252Q^* mice. Boxes indicate the median and interquartile range. Each dot represents an individual mouse; \*\**P* < 0.01, two-tailed Welch’s t-test.

### Gross brain abnormalities in *Lmx1b^R252Q/R252Q^* mice

During necropsy, we also noticed gross morphological abnormalities in the brains of *Lmx1b^R252Q/R252Q^* mice. Compared with wild-type controls, mutant brains appeared smaller and showed a more rounded overall morphology (Fig 4). In dorsal views, the bilateral inferior colliculi were separated at the midline in mutant brains, whereas this region was continuous and well organized in wild-type controls (Fig 4A, B). Mutant brains also showed cerebellar hypoplasia, characterized by a reduced cerebellar size relative to the overall brain morphology (Fig 4C, D). In addition, the olfactory bulb region appeared slightly distorted in several mutant brains. It remains unclear whether this appearance reflects an intrinsic alteration of the olfactory bulbs or differences in overall brain or cranial morphology. Given the established role of *Lmx1b* in midbrain-hindbrain boundary formation, the separation of the inferior colliculi and cerebellar hypoplasia observed in mutant mice are consistent with altered *Lmx1b* function.

**Fig 4.**
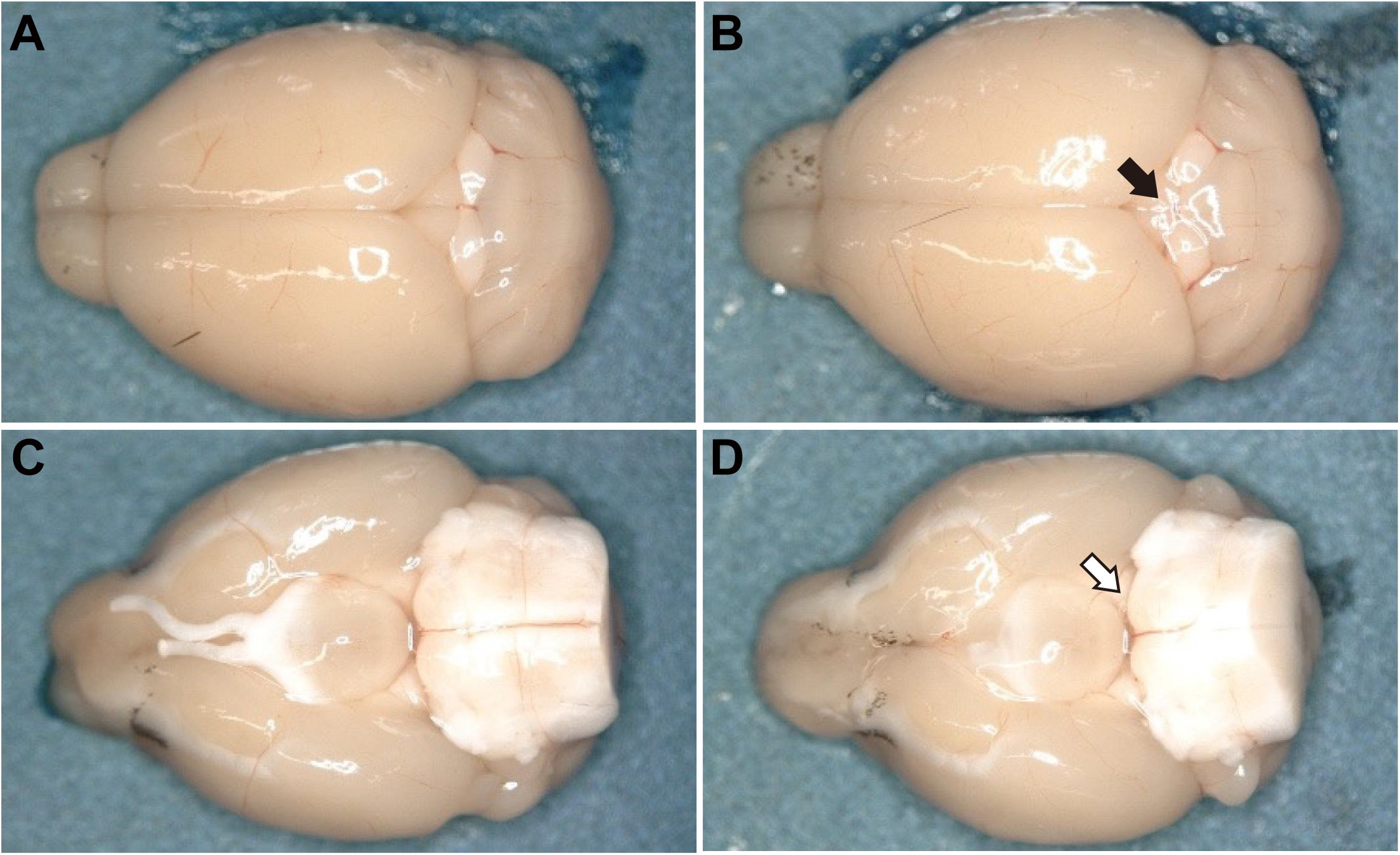
Gross brain abnormalities in *Lmx1b^R252Q/R252Q^* mice. (A, B) Dorsal views of whole brains from wild-type (A) and *Lmx1b^R252Q/R252Q^* mice (B). Mutant brains showed separation of the bilateral inferior colliculi at the midline, whereas this region was continuous in wild-type controls. An arrow indicates the affected region. (C, D) Ventral views showing cerebellar hypoplasia in mutant mice (an open arrow).

### *Lmx1b^R252Q/R252Q^* mice exhibit a relatively mild renal phenotype

To determine whether *Lmx1b^R252Q/R252Q^* mice recapitulate the nephropathy associated with human *LMX1B^R246Q^*, we examined their renal structure and function. Light microscopic examination revealed no overt renal abnormalities in mutant mice compared with wild-type controls (Fig 5A). HE staining showed preserved glomerular and tubular architecture, with no significant difference in the number of nuclei per glomerular cross-section. There was also no apparent inflammatory cell infiltration or other evidence of renal injury. PAS and MT staining showed no evidence of mesangial matrix expansion or interstitial fibrosis, respectively, while PAM staining revealed no overt abnormalities in glomerular basement membrane (GBM) architecture. To examine glomerular ultrastructure, we performed transmission electron microscopy of kidneys from *Lmx1b^R252Q/R252Q^* mice. In most regions examined, the GBM appeared continuous and relatively uniform, and podocyte foot processes were generally well preserved (Fig 5B). However, occasional focal regions showed irregularity and heterogeneous organization of the GBM (Fig 5C). Thus, although overall glomerular ultrastructure was largely preserved, limited focal GBM abnormalities were detectable in *Lmx1b^R252Q/R252Q^* kidneys.

**Fig 5.**
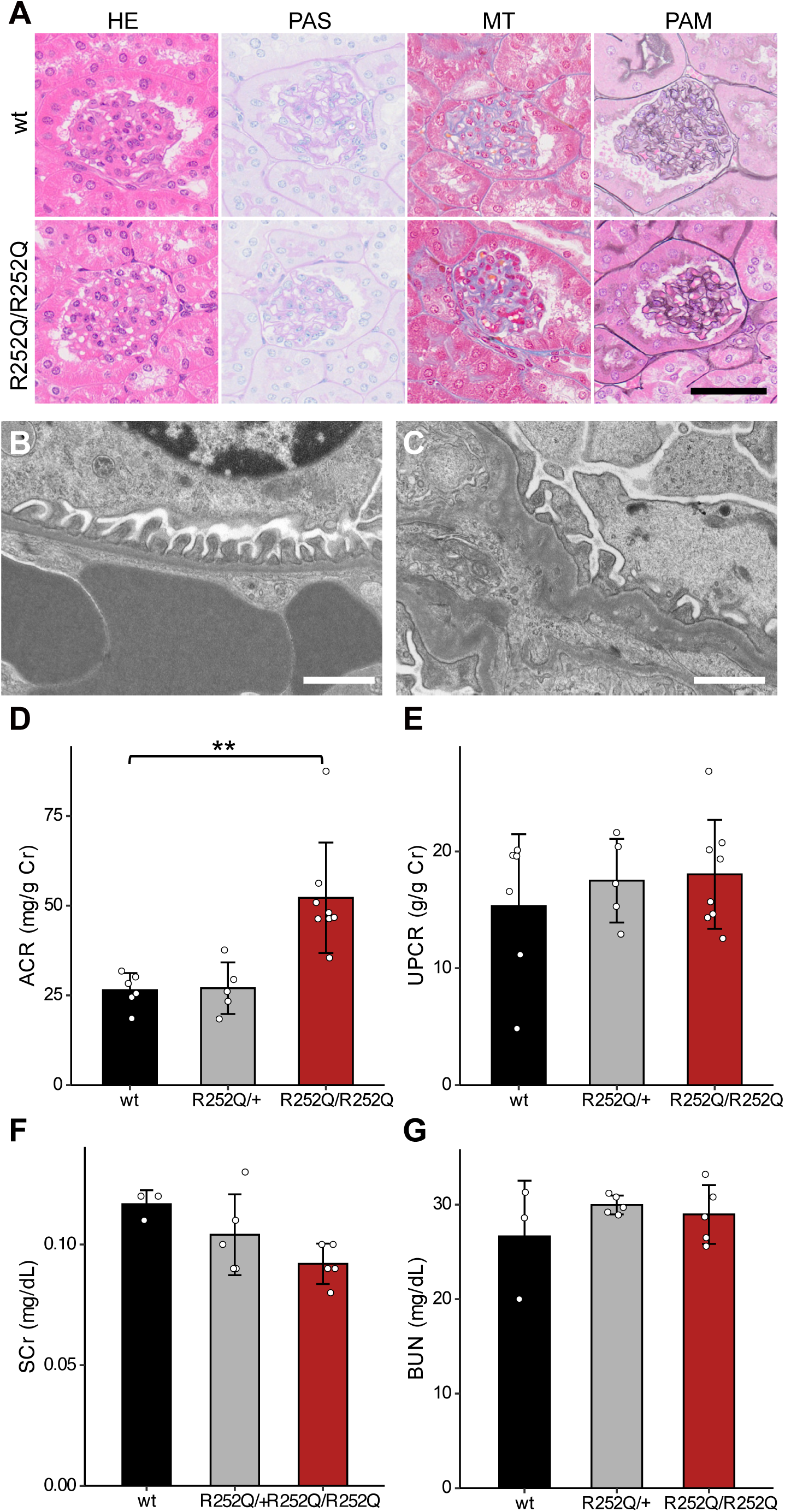
Mild renal involvement in *Lmx1b^R252Q/R252Q^* mice. (A) Representative kidney sections stained with HE, PAS, MT, and PAM, showing largely preserved glomerular structures in homozygous mutant mice. Scale bar, 50 µm. (B, C) Transmission electron microscopy of glomeruli from *Lmx1b^R252Q/R252Q^*mice showing largely preserved glomerular filtration barrier architecture in most areas (B) and focal GBM irregularity with podocyte foot process disorganization in selected regions (C). White scale bars, 1 µm. Urinary albumin-to-creatinine ratio (D) and urinary protein-to-creatinine ratio (E) in male wild-type, *Lmx1b^R252Q/+^*, and *Lmx1b^R252Q/R252Q^*mice. *Lmx1b^R252Q/R252Q^* mice showed increased urinary albumin excretion. Serum creatinine (F) and blood urea nitrogen (G) levels showed no significant differences among genotypes. (D-G) Data are shown as mean ± SD. Each dot represents an individual mouse. \*\**P* < 0.01, one-way ANOVA followed by Dunnett’s multiple-comparisons test, with wild-type mice as the control group.

To determine whether these structural findings were accompanied by renal functional abnormalities, urinary and serum biochemical parameters were measured in mutant and wild-type mice. The urinary albumin-to-creatinine ratio (ACR) was unchanged in *Lmx1b^R252Q/+^* mice but was significantly increased by approximately twofold in *Lmx1b^R252Q/R252Q^*mice compared with wild-type controls (Fig 5D). By contrast, the urinary protein-to-creatinine ratio (UPCR) did not differ significantly among the three genotypes (Fig 5E). Serum creatinine and blood urea nitrogen levels also did not differ significantly among the genotypes (Fig 5F, G). Other serum biochemical parameters, including total protein and albumin, showed no consistent genotype-dependent alterations (S1 Table). Thus, *Lmx1b^R252Q/R252Q^*mice exhibited a modest increase in urinary albumin excretion without a detectable increase in total urinary protein or overt impairment of renal function.

### Altered renal gene expression and reduced transcriptional activity of *Lmx1b^R252Q^*

To investigate molecular changes associated with the mild renal phenotype, we performed bulk RNA sequencing on whole-kidney RNA from male and female *Lmx1b^R252Q/R252Q^* and wild-type mice. Differential expression analysis identified 16 differentially expressed genes (DEGs) in males and 31 DEGs in females (Fig 6A, B, S2 Table; FDR < 0.05). Comparison of the two DEG sets identified four commonly upregulated genes (*Nell2*, *Gm31075*, *Smyd1*, and *Kirrel3*) and five commonly downregulated genes (*H2-Q6*, *H2-Q7*, *H2-Q8*, *Sema3g*, and *Shisa3*) in homozygous kidneys (Fig 6C). These shared changes included *Kirrel3* and *Sema3g*, which are associated with podocyte biology, as well as several non-classical MHC class I genes of the H2-Q family.

**Fig 6.**
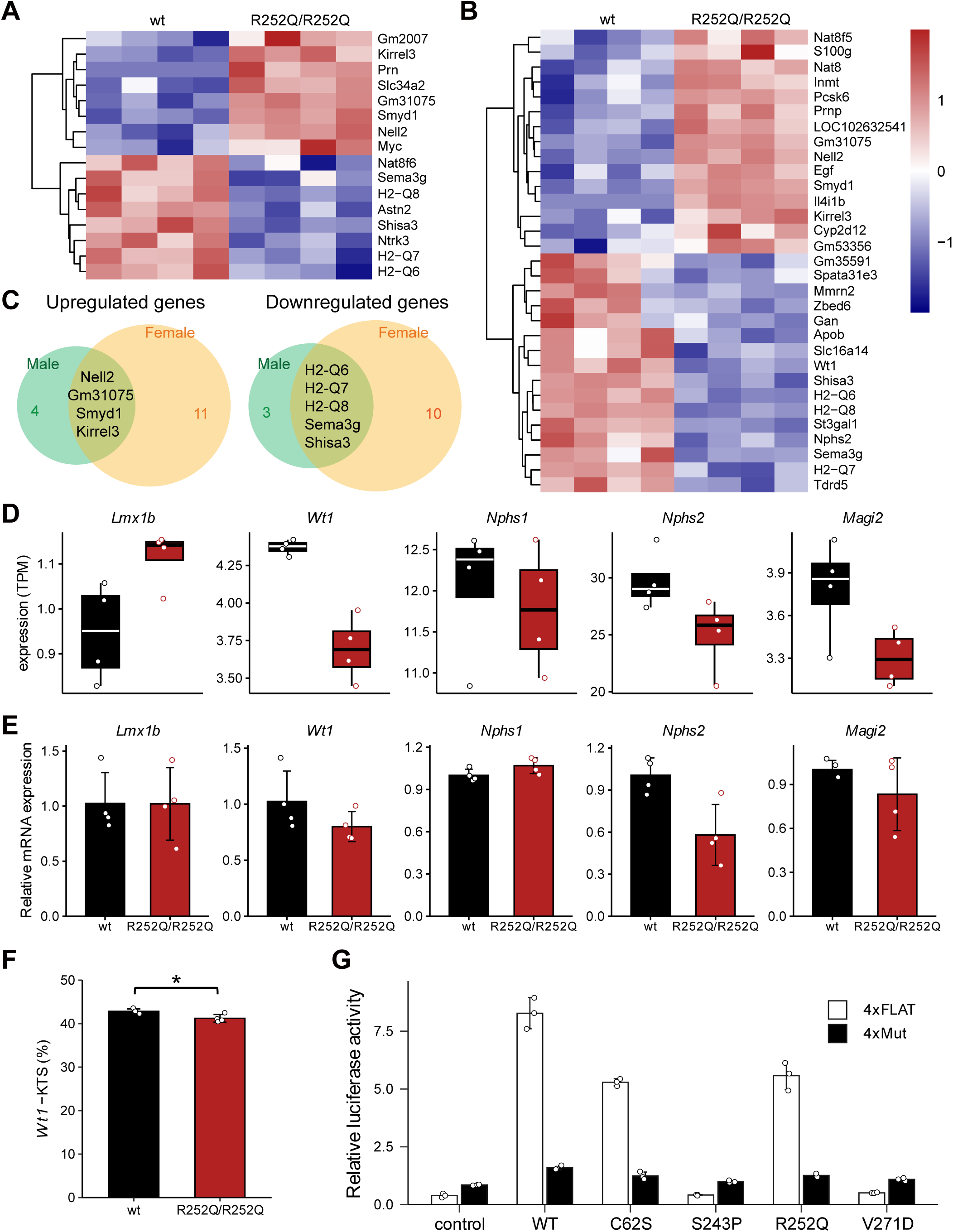
Renal transcriptional consequences of *Lmx1b^R252Q^* and functional characterization of *Lmx1b* variants. (A, B) Heatmaps of differentially expressed genes (DEGs) in the whole-kidney samples from male (A) and female (B) wild-type and *Lmx1b^R252Q/R252Q^*mice (n = 4 per genotype). DEGs were identified using DESeq2 with a false discovery rate (FDR) of < 0.05. Variance-stabilized expression values were standardized for each gene and are displayed as Z-scores, with red and blue indicating relatively high and low expression, respectively. (C) Overlap of significantly upregulated and downregulated genes between male and female kidneys. Genes within the overlapping regions indicate DEGs shared between the sexes. (D) RNA-seq derived TPM values for selected podocyte-associated genes in male kidneys. Black indicates wild-type mice and red indicates *Lmx1b^R252Q/R252Q^*mice. (E) Relative mRNA expression of podocyte-associated genes in male kidneys determined by quantitative PCR (n = 4 per genotype). Expression was normalized to *Atp5f1a* and is shown relative to the wild-type mean. Bars represent the mean ± SD, and dots represent individual mice. (F) Percentage of the *Wt1*(-KTS) isoform in male kidneys, determined by digital PCR. Bars represent the mean ± SD, and dots represent individual mice. (G) Transcriptional activities of wild-type LMX1B and the indicated mouse LMX1B variants assessed using luciferase reporters containing four tandem LMX1B-binding elements (4×FLAT) or mutated FLAT elements (4×Mut). An EGFP expression vector was used as the transfection control. Bars represent the mean ± SD, and dots represent individual measurements (n = 3). Statistical comparisons in (E) and (F) were performed using unpaired two-tailed Welch’s t-test. \**P* < 0.05. For the 4×FLAT reporter in (G), each variant was compared with WT using one-way ANOVA followed by Dunnett’s multiple-comparisons test. \*\**P* < 0.01.

In female homozygous kidneys, *Wt1* and *Nphs2*, both known downstream targets of LMX1B, were also identified among the downregulated genes (Fig 6B). We therefore examined the expression of these and other LMX1B-regulated podocyte genes in male kidneys. RNA-seq derived transcripts per million (TPM) values showed trends toward lower expression of *Wt1*, *Nphs2*, and *Magi2* in homozygous males (Fig 6D). Quantitative RT-PCR confirmed a significant reduction in *Nphs2* expression, whereas *Wt1* expression showed a nonsignificant downward trend (Fig 6E).

A previous study reported that the human LMX1B^R246Q^ variant preferentially reduced expression of the *WT1*(-KTS) isoform in cultured human podocytes [15]. We therefore examined whether the corresponding R252Q substitution affected *Wt1* isoform expression *in vivo*. Digital PCR analysis of the alternatively spliced *Wt1* isoforms revealed a significant reduction in the -KTS isoform in homozygous male kidneys (Fig 6F). Thus, although *Wt1* was not identified as a DEG in the male RNA-seq analysis, more detailed analyses demonstrated altered expression of LMX1B downstream targets in male homozygous kidneys.

The reduced expression of *Nphs2* and other podocyte-associated genes suggested that the R252Q substitution may impair LMX1B-mediated transcriptional regulation. We examined the transcriptional activity of mouse LMX1B^R252Q^ using a luciferase reporter containing four tandem LMX1B-binding motifs. Its activity was compared with that of selected patient-associated *LMX1B* variants and the mouse *Icst* mutant. Wild-type LMX1B strongly activated the 4×FLAT reporter, whereas the mutant reporter (4×Mut) showed only low activity across all conditions (Fig 6G). The R252Q variant showed reduced but clearly retained transcriptional activity compared with wild-type LMX1B. A similar partial reduction was observed for C62S, whereas S243P and V271D showed markedly impaired reporter activation. These results indicate that the R252Q variant causes a partial loss of LMX1B-mediated transcriptional activation rather than a complete loss of function.

## Discussion

We generated a knock-in mouse model carrying the *Lmx1b^R252Q^*variant corresponding to the recurrent human *LMX1B^R246Q^* variant associated with renal-predominant *LMX1B* disease. Homozygous *Lmx1b*^R252Q^ mice are viable and fertile but exhibit growth retardation, ocular opacity accompanied by glaucoma-like abnormalities, patella hypoplasia, and gross brain abnormalities. In contrast, their renal phenotype was relatively mild. Routine light microscopy revealed no overt lesions, whereas transmission electron microscopy detected occasional focal GBM abnormalities, and urinary analysis demonstrated modest albuminuria without increased total urinary protein or impaired renal function. These findings indicate that R252Q is a partially functional allele with tissue-dependent phenotypic consequences. The model reproduces several developmental manifestations of NPS but only partially recapitulates the renal-predominant phenotype associated with human R246Q.

Previous genotype-phenotype analysis showed that patients with *LMX1B* variants in the homeodomain had more frequent and greater proteinuria than those with variants in the LIM domains, suggesting that homeodomain dysfunction increases susceptibility to renal involvement, although variant location alone does not determine the phenotype [1]. Several variants associated with renal-predominant disease have been identified around Arg246. R246Q and R246P were reported in families with hereditary FSGS without the characteristic extrarenal manifestations of NPS [17], and additional patients and families carrying R246Q or R246L have subsequently been reported [16, 18]. A renal-predominant R249Q variant has also been identified at a nearby residue within helix II of the homeodomain [19]. Substitutions at or near Arg246 may have distinctive effects on LMX1B function. Isojima and colleagues demonstrated that human R246Q retained residual transcriptional activity despite a significant reduction relative to wild-type LMX1B. In the present study, the corresponding mouse R252Q substitution also retained partial activity in the reporter assay. Thus, R246Q/R252Q appears to represent a partial loss-of-function variant rather than a simple null allele. However, whether the renal-predominant phenotype results solely from a quantitative reduction in transcriptional activity or also involves qualitative changes in target-gene regulation remains unresolved.

A more severe form of homeodomain dysfunction is illustrated by the mouse *Lmx1b^Icst^* allele, which encodes the V271D substitution in the DNA-recognition helix. V271D abolished binding to the FLAT element and reduced reporter activity to background levels [13]. Although *Lmx1b^Icst/Icst^* mice exhibited a phenotype comparable to that of *Lmx1b* knockout mice, *Icst* is not a simple null allele. *Lmx1b^Icst/+^* mice exhibited glaucoma-like eye abnormalities and GBM abnormalities of varying severity, with partial postnatal lethality. The mutant protein retained the ability to participate in LDB1-mediated complexes with wild-type LMX1B, thereby reducing the abundance of functional complexes and producing a dominant-negative effect [13]. Unlike V271D, R252Q retained partial activity, and even homozygous mice showed only modest albuminuria and limited renal pathology. These observations indicate a relationship between the degree of homeodomain dysfunction and renal severity in mice. Because the distribution of organ involvement differs between human and mouse, the phenotypic gradient of LMX1B dysfunction is unlikely to be determined solely by residual transcriptional activity and may also reflect species-dependent differences in tissue-specific functional thresholds, target-gene regulation, or genetic modifiers.

The extrarenal abnormalities in *Lmx1b^R252Q/R252Q^* mice are consistent with the developmental functions of LMX1B. Patellar hypoplasia is compatible with its essential role in dorsal limb patterning, whereas the ocular abnormalities are consistent with previous mouse studies showing that *Lmx1b* is required for anterior segment development, trabecular meshwork formation, and maintenance of corneal transparency [12, 20]. Cerebellar hypoplasia and abnormalities around the midbrain–hindbrain region are also consistent with the established role of *Lmx1b* in midbrain and hindbrain development [11]. Thus, although R252Q retains transcriptional activity, this residual activity is insufficient to support normal development in several tissues when present in the homozygous state. The differences in severity among the skeletal, ocular, neural, and renal phenotypes suggest that the functional requirement for LMX1B differs among organs and developmental stages.

Despite the severe extrarenal defects, the renal phenotype of the *Lmx1b^R252Q^* mice was comparatively mild. Renal involvement in NPS is highly variable and often initially presents as proteinuria, while only a minority of affected individuals progress to end-stage kidney disease. Moreover, NPS-associated renal lesions require electron microscopy to reveal characteristic irregular thickening of the GBM and collagen fibril deposition [21–23]. The renal phenotype of *Lmx1b^R252Q/R252Q^*mice may therefore represent an early or subclinical alteration of the glomerular filtration barrier.

LMX1B maintains podocyte differentiation and function by regulating genes involved in the slit diaphragm and GBM, including *Col4a3*, *Col4a4*, *Nphs2*, and *Cd2ap* [8, 9, 24, 25]. A previous study further reported that human R246Q altered the expression of *WT1*, *NPHS1*, *TRPC6*, and *GLEPP1* in cultured podocytes and preferentially reduced *WT1*(-KTS) isoforms [15]. In the present mouse model, *Wt1*(-KTS) expression was also reduced, while *Nphs2* showed a significant decrease by qPCR. These findings suggest modest and selective changes in podocyte-associated gene expression, consistent with the small number of DEGs and mild renal phenotype. Although cell-type-specific changes may be underestimated by whole-kidney bulk RNA-seq, *Sema3g* and *Ntrk3* were downregulated, whereas *Kirrel3* was upregulated. These genes are associated with podocyte protection, cytoskeletal maintenance, and slit diaphragm function, respectively [26–28]. Together with the reductions in *Nphs2* and *Wt1*(-KTS), these changes suggest an alteration in pathways involved in podocyte maintenance and function. The reductions in several *H2-Q* family genes may also reflect subtle alterations in renal immune-related pathways, although their biological significance remains unclear in the absence of histological evidence of inflammatory infiltration.

Several limitations should be considered when interpreting this model. Human R246Q-associated disease occurs in the heterozygous state, whereas the clearest phenotypes in the present study were observed in homozygous mice. Heterozygous mice showed no significant increase in albuminuria. The renal analyses were also conducted under baseline conditions and at a limited age, and the effects of aging, genetic background, and secondary renal stress remain unknown. Longitudinal urinary analysis, aged cohorts, and renal challenge experiments may determine whether R252Q confers latent susceptibility that becomes apparent over time or after additional stress. Despite these limitations, the model combines clear NPS-related developmental abnormalities with modest albuminuria and limited renal pathology. It therefore provides a useful system for investigating the pleiotropic consequences of a partially functional LMX1B allele and the factors that modify renal susceptibility.

## Materials and Methods

### Breeding and genotyping of mice

C57BL/6N mice were purchased from CLEA Japan Inc. (Tokyo, Japan) and maintained under specific pathogen-free conditions at the RIKEN BioResource Research Center (BRC) in Tsukuba. *Lmx1b^R252Q^* mutant mice were maintained by backcrossing to C57BL/6N mice and genotyped at each generation. Experimental animals were obtained by intercrossing heterozygous mice. Genomic DNA was extracted from ear biopsies after overnight incubation at 55°C in lysis buffer (50 mM Tris-HCl pH 8.0, 100 mM EDTA, 100 mM NaCl, 1% SDS, and 0.5 mg/mL Proteinase K). Lysates were diluted and used directly as templates for allele specific PCR using the primers listed in S3 Table. Genotypes were determined based on the resulting amplification patterns. All animal experiments were conducted in accordance with the guidelines of the Institutional Animal Care and Use Committee of RIKEN Tsukuba Branch (Approval number: T2024-EP008).

### Genome editing

We introduced a patient-associated variant into the mouse *Lmx1b* locus using CRISPR/Cas9 genome editing. The mixture of gRNA, a single-stranded oligonucleotide (ssODN) donor (IDT, IA, USA), and recombinant Cas9 protein (IDT) was dissolved in Opti-MEM (Thermo Fisher Scientific, MA, USA) and introduced into fertilized eggs by electroporation. The sequences of the gRNA and ssODN donor used are listed in S3 Table. Fertilized mouse zygotes were obtained from superovulated female C57BL/6N mice. In brief, the female mice were injected intraperitoneally with 5 IU of pregnant mare serum gonadotropin (PMSG), followed 48 hours later by 5 IU of human chorionic gonadotropin (hCG), and subsequently mated with male mice. Fertilized one-cell embryos were collected and electroporated with gRNA/ssODN/Cas9 mixtures using a NEPA 21 electroporator (NEPAGENE Co. Ltd., Chiba, Japan). The electroporated embryos that developed to the two-cell stage were transferred to the oviducts of pseudopregnant ICR female mice to obtain founder mice. The targeted sequence was confirmed by Sanger sequencing during establishment of the mouse line. The resulting mouse strain, C57BL/6N-*Lmx1b^em1Amano^* (RBRC12972), is available from the RIKEN BRC.

### RNA extraction and expression analysis

Total RNA was extracted from whole kidneys with TRIzol reagent (Thermo Fisher Scientific). Library preparation and sequencing were performed by Macrogen Japan (Tokyo, Japan). Libraries were constructed using the TruSeq Stranded mRNA kit (Illumina, CA, USA) and sequenced on an Illumina NovaSeq X platform to generate paired-end 100 bp reads. The resulting FASTQ files were imported to the CLC Genomics Workbench v.26.0.2 (QIAGEN, Hilden, Germany) and mapped to the mouse reference genome (GRCm39/mm39). Read counts and TPM values were calculated and used for downstream analysis using R. Differential gene expression was evaluated using DESeq2 in R with significance defined as false discovery rate (FDR) < 0.05 [29]. The RNA-seq data generated in this study have been deposited in the DDBJ Sequence Read Archive under accession number PRJDB42757.

Quantitative RT-PCR analysis was performed using the One Step TB Green PrimeScript PLUS RT-PCR Kit (Takara Bio, Shiga, Japan) on a LightCycler 96 system (Roche Diagnostics GmbH, Mannheim, Germany) according to the manufacturer’s instructions. Relative gene expression levels were calculated using the ΔΔCq method, with *Atp5f1a* used as an internal control. Primers were designed using GETPrime [30], and the sequences were listed in S3 Table. The relative abundance of the *Wt1*(+KTS) and *Wt1*(-KTS) splice isoforms was determined by digital PCR using a custom TaqMan SNP Genotyping Assay on a QuantStudio Absolute Q Digital PCR System (Thermo Fisher Scientific). The proportion of each isoform was calculated from the concentrations measured using the corresponding FAM- and VIC-labeled probes.

### Urine and serum collection

Urine samples were collected by gently restraining mice and retrieving spontaneously voided urine from a clean collection plate. Urinary concentrations of albumin, creatinine, and total protein were measured by Oriental Yeast Co., Ltd. (Shiga, Japan) using standard methods. Blood samples were obtained from the orbital venous plexus under isoflurane anesthesia using a Pasteur pipette. Blood was kept at room temperature for 1 hour and then centrifuged at 1,700 × g for 15 minutes to obtain serum. Serum biochemical parameters were analyzed by the Japan Mouse Clinic of the RIKEN BRC.

### Histopathological and skeletal morphology analysis

Eyes were fixed overnight at room temperature in modified Davidson’s fixative, washed with 70% ethanol, and embedded in paraffin. Eye sections were stained with hematoxylin and eosin (HE) at the Japan Mouse Clinic of the RIKEN BRC. Kidneys for light microscopic analysis were fixed overnight at room temperature in Bouin’s fixative, washed with 70% ethanol, and embedded in paraffin. Kidney sections were stained with HE, periodic acid Schiff (PAS), periodic acid methenamine silver (PAM), and Masson’s trichrome (MT) by Morphotechnology Co., Ltd. (Hokkaido, Japan). For transmission electron microscopy (TEM) observation, kidneys were fixed in phosphate-buffered glutaraldehyde solution overnight at 4°C. The fixed samples were processed and analyzed by Hanaichi UltraStructure Research Institute, Co., Ltd. (Aichi, Japan). Skeletal morphology of adult hindlimbs was evaluated by X-ray micro-computed tomography (µCT) at the Japan Mouse Clinic of the RIKEN BRC. After fixation, hindlimb samples were stored in 70% ethanol until imaging. µCT imaging was performed using the ScanXmate-E090S system (Comscantecno Co., Ltd., Kanagawa, Japan) at an accelerating voltage of 40 kV and an X-ray tube current of 100 μA, with a 360° rotation, 1,200 projections, and an isotropic voxel size of 19.209 μm. Projection data were reconstructed into three-dimensional datasets using the filtered back-projection algorithm. The patella was visualized using OsiriX MD (Pixmeo SARL, Geneva, Switzerland), and patellar long-axis length was measured. The reconstructed datasets were segmented using Amira 2024.1 (Thermo Fisher Scientific), and patellar volume was determined from the segmented structures using the same software.

### Luciferase assay

The protein-coding sequence of mouse *Lmx1b* was amplified by PCR and cloned into pcDNA3.1 (Thermo Fisher Scientific). Based on this plasmid, pathogenic variants, including those identified in patients and mice, were introduced by site-directed mutagenesis. Sense and antisense oligonucleotides containing four tandem copies of the FLAT-like element identified in the promoter region of the human *NPHS2* gene [8] were annealed and inserted upstream of a minimal promoter and firefly luciferase in the plasmid pGL4.23 (Promega Corporation, WI, USA). The plasmid pGL4.74 ubiquitously expressing *Renilla* luciferase was used as an internal control. NIH3T3 cells (RCB2767) were obtained from the RIKEN BRC Cell Bank and maintained in Dulbecco’s modified Eagle’s medium containing 10% fetal bovine serum and 100 µg/mL penicillin-streptomycin. Cells at 70-90% confluency were transfected with firefly and *Renilla* luciferase reporter plasmids and *Lmx1b*-expressing plasmids using Lipofectamine 3000 (Thermo Fisher Scientific). Luciferase activity in the cell lysates was measured using the Dual-Luciferase Reporter Assay System (Promega Corporation) according to the manufacturer’s protocol.

### Statistical analysis

Statistical analyses were performed using R (version 4.3.3). Comparisons between two groups were performed using an unpaired two-tailed Welch’s t-test. Comparisons among three or more groups were performed using one-way analysis of variance (ANOVA) followed by Dunnett’s multiple-comparisons test. For quantitative PCR analyses, statistical tests were performed using ΔCq values. Individual data points are shown in the figures, and the statistical tests used are specified in the corresponding figure legends.

## Acknowledgments

We thank the animal facility staff at RIKEN BRC for animal care, the RIKEN BRC Japan Mouse Clinic staff for technical assistance with blood biochemical analyses, and Ayaka Saito for assistance with plasmid construction.

## Supporting information captions

**S1 Table. Blood biochemical parameters in *Lmx1b^R252Q^*mice.**

**S2 Table. DESeq2 results from kidney RNA-seq analyses in male and female mice.**

**S3 Table. Oligonucleotides used in this study.**

## References

1. Bongers EMHF, Huysmans FT, Levtchenko E, De Rooy JW, Blickman JG, Admiraal RJC, et al. Genotype– phenotype studies in nail-patella syndrome show that LMX1B mutation location is involved in the risk of developing nephropathy. European Journal of Human Genetics. 2005;13(8):935–46.

2. Ghoumid J, Petit F, Holder-Espinasse M, Jourdain AS, Guerra J, Dieux-Coeslier A, et al. Nail-Patella Syndrome: clinical and molecular data in 55 families raising the hypothesis of a genetic heterogeneity. Eur J Hum Genet. 2016;24(1):44–50.

3. Sweeney E, Fryer A, Mountford R, Green A, McIntosh I. Nail patella syndrome: a review of the phenotype aided by developmental biology. J Med Genet. 2003;40(3):153–62.

4. Castilla-Ibeas A, Zdral S, Oberg KC, Ros MA. The limb dorsoventral axis: Lmx1b’s role in development, pathology, evolution, and regeneration. Developmental Dynamics. 2024;253(9):798–814.

5. Chen H, Lun Y, Ovchinnikov D, Kokubo H, Oberg KC, Pepicelli CV, et al. Limb and kidney defects in Lmx1b mutant mice suggest an involvement of LMX1B in human nail patella syndrome. Nature Genetics. 1998;19(1):51–5.

6. Stenson PD, Mort M, Ball EV, Chapman M, Evans K, Azevedo L, et al. The Human Gene Mutation Database (HGMD®): optimizing its use in a clinical diagnostic or research setting. Human Genetics. 2020;139(10):1197– 207.

7. Landrum MJ, Lee JM, Benson M, Brown GR, Chao C, Chitipiralla S, et al. ClinVar: improving access to variant interpretations and supporting evidence. Nucleic Acids Research. 2018;46(D1):D1062–D7.

8. Miner JH, Morello R, Andrews KL, Li C, Antignac C, Shaw AS, et al. Transcriptional induction of slit diaphragm genes by Lmx1b is required in podocyte differentiation. J Clin Invest. 2002;109(8):1065–72.

9. Rohr C, Prestel J, Heidet L, Hosser H, Kriz W, Johnson RL, et al. The LIM-homeodomain transcription factor Lmx1b plays a crucial role in podocytes. Journal of Clinical Investigation. 2002;109(8):1073–82.

10. Kishio N, Iwama K, Nakanishi S, Shindo R, Yasui M, Nicho N, et al. A deletion variant in LMX1B causing nail–patella syndrome in Japanese twins. Human Genome Variation. 2024;11:10.

11. Guo C, Qiu HY, Huang Y, Chen H, Yang RQ, Chen SD, et al. Lmx1b is essential for Fgf8 and Wnt1 expression in the isthmic organizer during tectum and cerebellum development in mice. Development. 2007;134(2):317–325.

12. Pressman CL, Chen H, Johnson RL. lmx1b, a LIM homeodomain class transcription factor, is necessary for normal development of multiple tissues in the anterior segment of the murine eye. genesis. 2000;26(1):15–25.

13. Cross SH, Macalinao DG, McKie L, Rose L, Kearney AL, Rainger J, et al. A dominant-negative mutation of mouse Lmx1b causes glaucoma and is semi-lethal via LDB1-mediated dimerization [corrected]. PLoS Genet. 2014;10(5):e1004359.

14. Konomoto T, Imamura H, Orita M, Tanaka E, Moritake H, Sato Y, et al. Clinical and histological findings of autosomal dominant renal-limited disease with LMX1B mutation. Nephrology (Carlton). 2016;21(9):765–773.

15. Hall G, Lane B, Chryst-Ladd M, Wu G, Lin JJ, Qin X, et al. Dysregulation of WTI (-KTS) is Associated with the Kidney-Specific Effects of the LMX1B R246Q Mutation. Sci Rep. 2017;7:39933.

16. Isojima T, Harita Y, Furuyama M, Sugawara N, Ishizuka K, Horita S, et al. LMX1B mutation with residual transcriptional activity as a cause of isolated glomerulopathy. Nephrology Dialysis Transplantation. 2014;29(1):81–8.

17. Boyer O, Woerner S, Yang F, Oakeley EJ, Linghu B, Gribouval O, et al. LMX1B mutations cause hereditary FSGS without extrarenal involvement. J Am Soc Nephrol. 2013;24(8):1216–22.

18. Li X, Fan J, Fu R, Peng M, He J, Chen Q, et al. Case report: A novel R246L mutation in the LMX1B homeodomain causes isolated nephropathy in a large Chinese family. Medicine (Baltimore). 2024;103(10):e37442.

19. Edwards N, Rice SJ, Raman S, Hynes AM, Srivastava S, Moore I, et al. A novel LMX1B mutation in a family with end-stage renal disease of ’unknown cause’. Clinical Kidney Journal. 2015;8(1):113–9.

20. Liu P, Johnson RL. Lmx1b is required for murine trabecular meshwork formation and for maintenance of corneal transparency. Developmental Dynamics. 2010;239(8):2161–71.

21. Hoyer JR, Michael AF, Vernier RL. Renal disease in nail-patella syndrome: clinical and morphologic studies. Kidney Int. 1972;2(4):231–8.

22. Taguchi T, Takebayashi S, Nishimura M, Tsuru N. Nephropathy of nail-patella syndrome. Ultrastruct Pathol. 1988;12(2):175–83.

23. Najafian B, Smith K, Lusco MA, Alpers CE, Fogo AB. AJKD Atlas of Renal Pathology: Nail-Patella Syndrome–Associated Nephropathy. American Journal of Kidney Diseases. 2017;70(4):e19–e20.

24. Burghardt T, Kastner J, Suleiman H, Rivera-Milla E, Stepanova N, Lottaz C, et al. LMX1B is essential for the maintenance of differentiated podocytes in adult kidneys. J Am Soc Nephrol. 2013;24(11):1830–48.

25. Morello R, Zhou G, Dreyer SD, Harvey SJ, Ninomiya Y, Thorner PS, et al. Regulation of glomerular basement membrane collagen expression by LMX1B contributes to renal disease in nail patella syndrome. Nature Genetics. 2001;27(2):205–8.

26. Ishibashi R, Takemoto M, Akimoto Y, Ishikawa T, He P, Maezawa Y, et al. A novel podocyte gene, semaphorin 3G, protects glomerular podocyte from lipopolysaccharide-induced inflammation. Scientific Reports. 2016;6(1):25955.

27. Gerke P, Sellin L, Kretz O, Petraschka D, Zentgraf H, Benzing T, et al. NEPH2 is located at the glomerular slit diaphragm, interacts with nephrin and is cleaved from podocytes by metalloproteinases. J Am Soc Nephrol. 2005;16(6):1693–702.

28. Lepa C, Hoppe S, Stober A, Skryabin BV, Sievers LK, Heitplatz B, et al. TrkC Is Essential for Nephron Function and Trans-Activates Igf1R Signaling. J Am Soc Nephrol. 2021;32(2):357–74.

29. Love MI, Huber W, Anders S. Moderated estimation of fold change and dispersion for RNA-seq data with DESeq2. Genome Biology. 2014;15:550.

30. David FPA, Rougemont J, Deplancke B. GETPrime 2.0: gene- and transcript-specific qPCR primers for 13 species including polymorphisms. Nucleic Acids Research. 2017;45(D1):D56–D60.

